# X-ray induced bio-acoustic emissions from cultured cells

**DOI:** 10.1101/2021.06.14.448285

**Authors:** Bruno F.E. Matarèse, Hassan Rahmoune, Nguyen T. K. Vo, Colin B. Seymour, Paul N. Schofield, Carmel Mothersill

## Abstract

**Purpose:** We characterise for the first time the emission of bio-acoustic waves from cultured cells irradiated at doses of X-ray photon radiation relevant to medical and accidental exposure.

**Methods and materials:** Human cancer cell lines (MCF-7, HL-60) and control cell-free media were exposed to 1 Gy X-ray photons while recording the sound generated before, during and after irradiation. Cellular cytotoxicity following photon irradiation was determined by extracellular LDH levels, and irradiated cell conditioned media were tested for their ability to elicit a bystander effect in reporter cells.

**Results:** We report the first recorded acoustic signals captured from a collective pressure wave response to ionising irradiation. The signature of the collective acoustic peaks was temporally wider and with higher acoustic power for irradiated HL-60 than for irradiated MCF-7.

**Conclusions:** We show that at doses of X-ray irradiation capable of producing the release of bystander effect-inducing activity, both cell types emit a characteristic acoustic signal for the duration of the radiation pulse. The rapid signal decay is consistent with a passive rather than an active acoustic signal generation. This preliminary study suggests that further work on the potential role of radiation induced acoustic emission (RIAE) in the cellular bystander effect is merited.

## Introduction

Intercellular and inter-organismal communication of an “irradiated state” constituting the radiation-induced bystander effect (RIBE) remains to be a widely researched topic with many unknowns (Mothersill and Seymour 2019). How RIBEs are initiated and transmitted, especially at low doses, still needs to be realized for a fuller picture of biological events taking place at the molecular and sub-atomic levels. A cumulative body of evidence suggests that in addition to biochemical factors mediating RIBEs (Azzam et al. 2001, Le et al. 2017, Mothersill and Seymour 1998, Mothersill, Smith, et al. 2007), biophysical elements can contribute to RIBEs (Cohen et al. 2020, Le et al. 2015, Mothersill et al. 2014, Mothersill, Moran, et al. 2007, Mothersill et al. 2012). The possibility that radiation-induced bio-acoustic signals might be involved in eliciting RIBEs was previously suggested, though not experimentally verified, by Mothersill et al. (2011), who found that signalling between irradiated and unirradiated fish occurred even when water-borne diffusion or light photon signalling was blocked. In this preliminary study we explore the generation of an acoustic signal by irradiated cells.

There is an intrinsic relationship between electromagnetic forces (mediated by photons) and acoustic vibration (mediated by phonons), experimentally established in the 19^th^ century (Bell 1881). On exposure to ionising photons, inner-shell electrons are excited, generating photoelectrons, Auger electrons, or electromagnetic radiation (e.g. photoluminescence), which decay producing cascades of secondary electrons as these decay processes transfer kinetic energy to surrounding atoms to reach thermal equilibrium. These processes lead to a transient thermoelastic expansion of the biological structure generating the pressure waves of an acoustic emission (Garcia et al. 1988)

We recently proposed the hypothesis that acoustic energy released on interactions of biota with electromagnetic radiation may represent a signal released by irradiated cells leading to, or complementing, or interacting with, other responses, such as endosome release, responsible for signal relay within the unirradiated individuals in the targeted population (Matarèse et al. 2020). In the current study, the primary objective was to experimentally demonstrate, for the first time, the bio-acoustic wave emissions generated as biological cellular response during X-ray photon irradiation of cells in culture. We provide the first recorded acoustic signals captured from a collective pressure wave response corresponding to the thermoelastic expansion of cells during low dose X-ray radiation. We demonstrate here that irradiation with 1Gy of 6 MeV X ray photons causes cultured cells to emit characteristic sound waves during irradiation and propose that this acoustic signal might trigger, or contribute to the triggering, of the bystander response in neighbouring cells.

## Materials and methods

### Cell culture and irradiation

Human leukaemia cells HL-60 and breast cancer cell line MCF-7 were cultured in RPMI and DMEM media (Sigma-Aldrich), respectively, in the presence of 10% (v/v) heat-inactivated foetal calf serum (Thermo Fisher Scientific), 50 U/mL penicillin, and 50 µg/mL streptomycin (Sigma). Cells were seeded on a gelatin matrix in 40-mm petri dishes (Sigma-Aldrich) and incubated at 37°C in 95% ambient air and 5% (v/v) CO_2_. HL-60 and MCF-7 cells seeded at 1.5 × 10^6^ cells per 4 cm diameter petri dish and exposed 24 hours later to 1 Gy of 6 MeV X-ray photons generated by a Varian Clinac 2100 at 0.37 Gy/minute. At exposure adherent cells were 80% confluent and viability as measured by trypan blue exclusion 95% and 90% for MCF-7 and HL-60 respectively.

Human colorectal carcinoma HCT116 p53^+/+^ cells, used to assay bystander activity, were cultured in RPMI medium supplemented with 10% (v/v) heat inactivated fetal bovine serum, 2 mM L-glutamine, 25 mM HEPES, 100 U/mL penicillin, and 100 µg/mL streptomycin (Sigma-Aldrich).

### Bio-acoustic signal recording

Acoustic emissions were measured using a custom large-bandwidth ultrasound transducer with a central frequency of 1 MHz placed in a 40-mm diameter petri dish containing the media inside. The transducer was immersed on the inner side of the petri dish and was not exposed to the X-ray irradiation area. The transducer was connected to a low-noise pre-amplifier with approximately 60 dB gain and recorded with a PicoScope oscilloscope at sample rate 9.766 MS/s with 102,4 ns sample interval. Signals were read out before, during and after dose delivery using cell-free culture media alone as control.

### Cell membrane integrity/cytotoxicity measurements – Lactate dehydrogenase (LDH) assay

An hour after exposure culture media were collected and cleared by centrifugation at 100 x g at 5°C for 5 min to remove any remnant of cells. The LDH assays were performed in duplicate with 50 μL conditioned cell culture media per well using the Abcam LDH (cytotoxicity) assay kit (#ab65393). The quantity of nicotinamide adenine dinucleotide (NADH) was assessed spectrophotometrically at 450 nm using the Tecan Infinite M200 Pro microplate reader with integrated command via the i-control® software and by mixing the NADH detection buffer with the supernatant according to the manufacturer’s instructions.

### Induction of mitochondrial membrane potential alterations

Cell culture conditioned medium (CCCM) and Irradiated cell culture conditioned medium (ICCM) were tested for their effects on the mitochondrial membrane potential of bystander reporter HCT116 p53^+/+^ cells. Briefly, reporter cells were exposed to CCCM or ICCM for 1 h at 37C. Then the reporter cells were incubated with the potentiometric fluorescent dye JC-1 to measure the relative mitochondrial membrane potential as previously described by Vo et al. (Vo et al. 2017). Relative fluorescent units were measured with the Tecan Infinite M200 Pro microplate reader with integrated command via the i-control^®^ software.

### Statistical Analysis

The bio-acoustic peaks observed in the recorded sound signals were analyzed using Matlab® software. The average temporal ‘‘width’’ and acoustic power ‘‘height’’ for each sound peak was calculated for the total peaks observed during a set period of time of one second and were carried out using n=5 biological replicates. The number of peaks per second over a sound pressure voltage over a threshold of -3.5V were determined for same n=5 replicates. LDH assays and Mitochondrial membrane potential were carried out using n=5 replicates. Significant differences between groups were determined using a two-tailed Mann-Whitney U test. Value of *p* <0.05 was deemed statistically significant.

## Results

### Bio-acoustic signals induced by X-Ray radiation

Sound wave signals were recorded before, during and after X-ray radiation in order to experimentally verify the bio-acoustic wave emission generated from a cell-cultured population. The recording of the raw temporal acoustic signal for adherent MCF-7; non-adherent HL-60 cells and respective media control RPMI or DMEM is represented in Figure 2. No sound peaks are observed for the control media samples while sharp sound pressure peaks are recorded for MCF-7 and HL-60 samples which represent the collective pressure response of the cell population in culture. We demonstrate that during the period of dose delivery, the non-adherent HL-60 cells generated spikes of a sharp acoustic signal of average temporal width of 20 μs at an average rate interval of 357Hz between each spike. Adherent MCF-7 cells generate spikes of a sharp acoustic signal of average temporal width of 18 μs at an average rate interval of 344Hz between each spike. The transient thermoelastic expansion of cells during X-ray irradiation generates collective pressure waves that alternate between pressurised and de-pressured phases with observed resting interval before each peak elicited. The sharp pressure spikes start with a pressurisation period and followed by de-pressurisation period.

**Figure 1.**
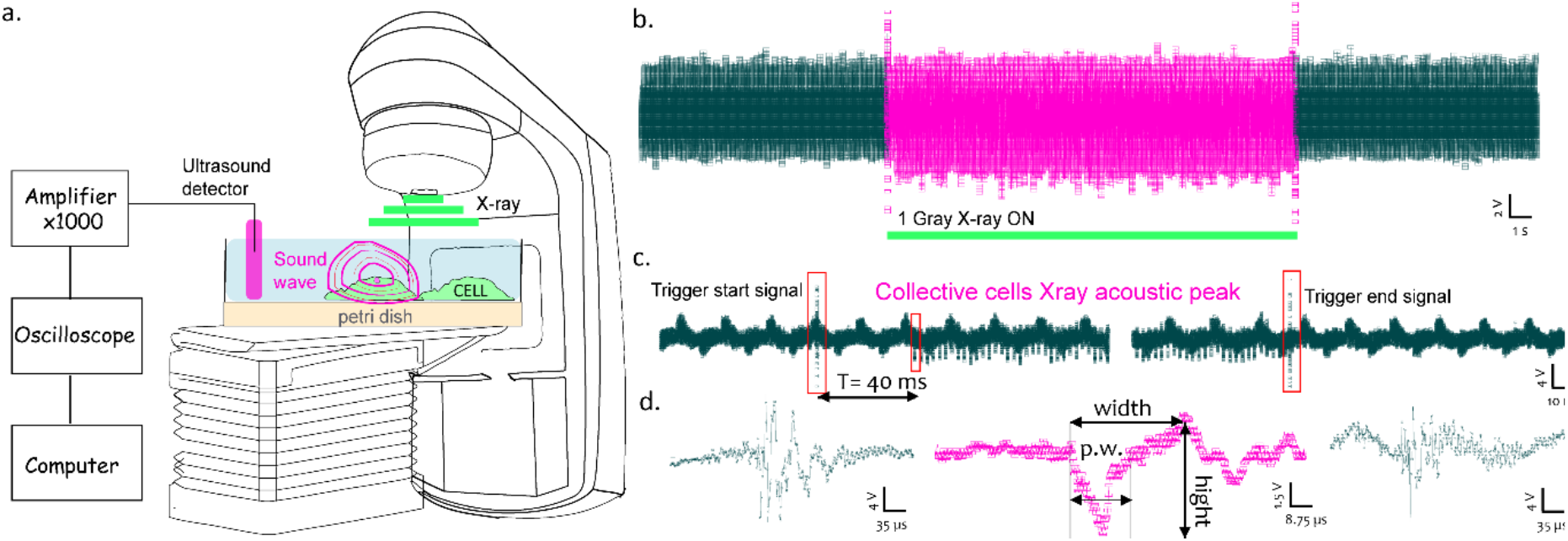
Combined X-ray radiation stimuli with sound recording setup and signal information. a.) schematic experimental setup showing amplified sound recording system combined with Varian Clinac arrangement with cultured cells. b.) Raw signal showing sound signal recorded before, during and after X-ray radiation. c.) Zoomed raw sound signal recorded at the time of X-ray triggered start & end signal observed and d.) Zoomed raw sound signal of X-ray acoustic triggered start peak (left in black); bio-acoustic cell population peak (middle in pink) and X-ray acoustic triggered end peak (right in black).

**Figure 2:**
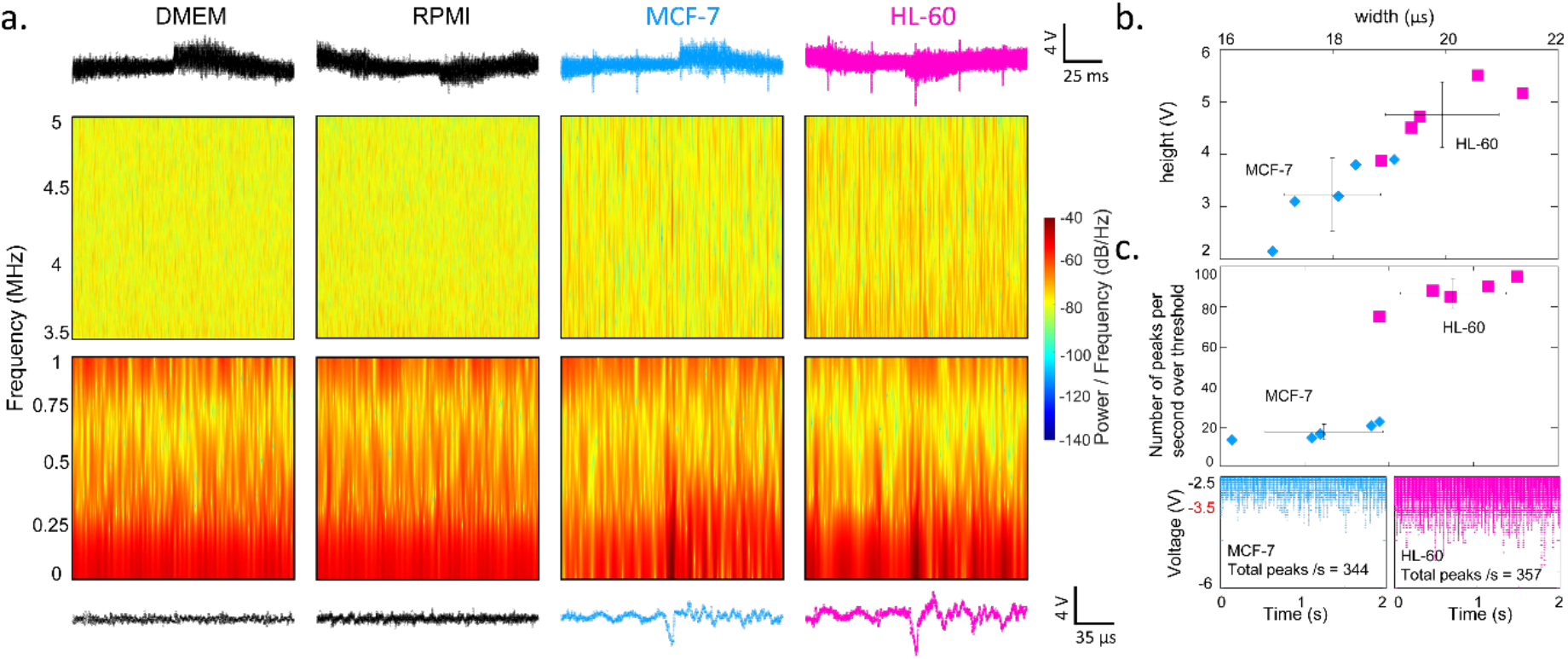
Bio-acoustic signal information. a.) Raw temporal acoustic signals for adherent MCF-7 (light blue); suspended HL-60 (pink) cells sample and respective media control RPMI or DMEM (black) and the respective frequency spectrum map of processed acoustic signals for low frequency [0 - 1MHz] and for high frequency [3.5 - 5MHz] domains as a function of irradiation time b.) Sound pressure peak analysis shown for the average temporal ‘‘width’’ (in μs) vs. the average sound power amplitude ‘‘height’’ (in mV) and c.) the average number of peaks per second over the electro-acoustic potential threshold of (−3.5V) for N=5 replicates and the overall average and standard deviations for adherent MCF-7 (light blue); suspended HL-60 (pink) cell-types.

The acoustic pressure peak analysis is based on temporal width (in Ds) and sound power amplitude by electro-acoustic signal height (in mV). The frequency spectrum map of processed acoustic signals for lower range of frequency [0 - 1MHz] and for high frequency [3.5 - 5MHz] domains as a function of irradiation time shows that the sharp spikes observed correspond to a lower frequency range from 10^−2^ Hz to 750 kHz and weaker ultrasound signals at higher frequencies observed between 3.5 MHz and 4.8 MHz.

The number of peaks per second over the electro-acoustic potential threshold of (−3.5V) further highlighted the different biosignature between the adherent and non-adherent cells with 15 peaks/s compared to 87 peaks/s respectively. The sound signal from MCF-7 cells showed a similar frequency range, but a weaker acoustic power was observed with electro-acoustic signal amplitude of 2 V decrease overall for each spike which can reflect the fact of having less number of cells per area. The signal does not continue after the end of irradiation, indicating relatively efficient relaxation processes.

### Cell damage and bystander factor production

In order to demonstrate that in this experiment irradiation caused cellular damage and induction of the release of bystander inducing factor as previously reported we examined both effects in the same cells from which acoustic signals were measured.

Both MCF-7 and HL-60 cells have been previously shown to produce a bystander effect (Al-Mayah et al. 2015, Ammar et al. 2012, Mohd Zainudin et al. 2020, Pandey et al. 2011b, Shao et al. 2008). HL-60 cells have also been shown to manifest an increase in intracellular reactive oxygen species (ROS) and alterations in mitochondrial membrane potential (Saenko et al. 2013). To assess immediate radiation damage we used LDH as a marker for membrane integrity (Niles et al. 2009, Uchide et al. 2009) and measured the generation of extracellular LDH (Figure 3) in both cell types. A significant (*p*=0.0158 two tailed Mann-Whitney test) increase in LDH was found in HL-60 irradiated cell conditioned medium (ICCM) while a small but not significant increase was found in that from MCF-7 cells. We therefore see a differential response to irradiation on extracellular levels of LDH, used as a cytotoxicity and/or membrane integrity status (Kumar et al. 2018) of non-adherent HL-60 and adherent MCF-7 cells. HL-60 seems to be more sensitive to irradiation as the extracellular level of LDH nearly doubled (from 3.4 +/-SE to 7.04 +/-SE) post exposure (Figure 3a). Such membrane fragility or cytotoxicity was not observed where similar exposure was performed on MCF-7 (Figure 3a). Increased cytotoxicity and membrane fragility seen in non-adherent HL-60 cells post-exposure may be reflected in a reduced mitochondrial membrane potential in bystander assay cells (HCT116 p53^+/+^) when challenged with ICCM from HL-60.

**Figure 3:**
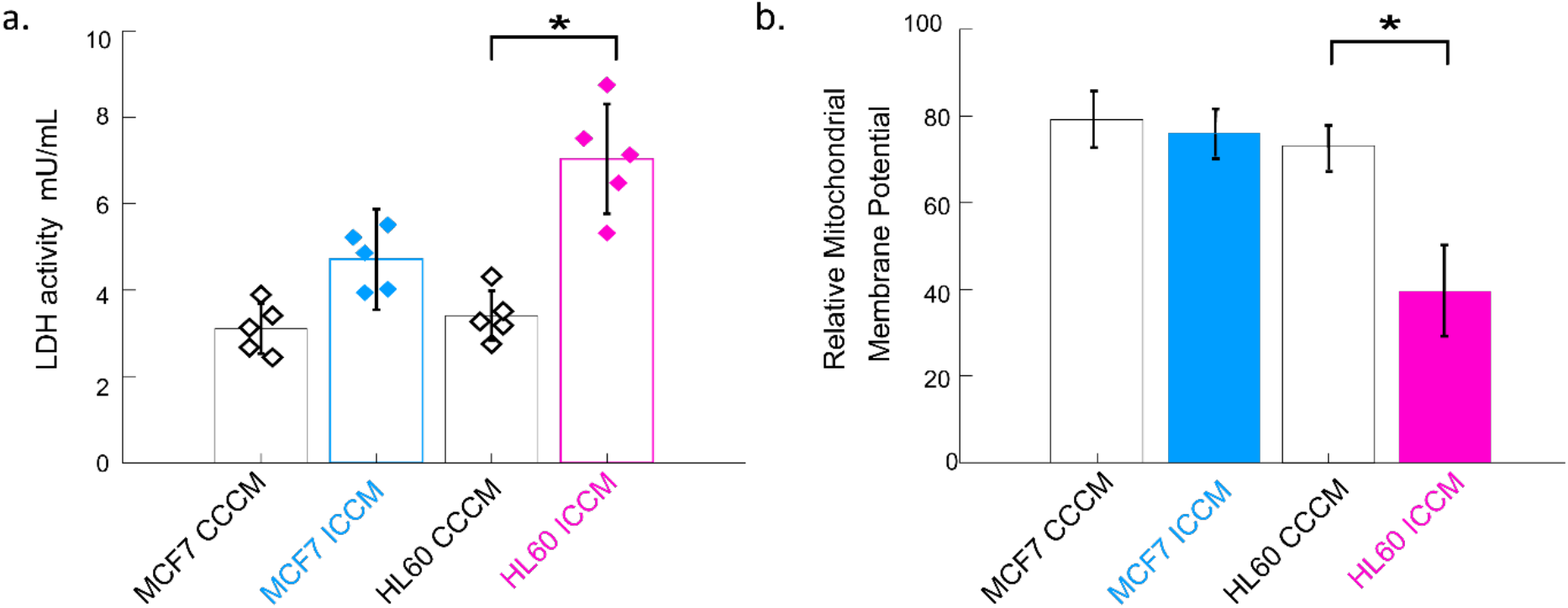
Cell membrane integrity; cytotoxicity and bystander effect. a.) MCF-7 and HL-60 extracellular LDH activity measurements b.) HCT116 cells mitochondrial membrane potential response post exposure to irradiated medium form HL-60 or MCF-7 cells in culture. RPMI or DMEM media was used as standalone control.

## Discussion

The observation that the ‘post-irradiated state”, comprising genomic instability and delayed cell death amongst other cell-type dependent phenomena, can be passed from an irradiated cell, tissue or whole organism to another, raises the important question of how the information might be transferred. Studies carried out by the authors (Mothersill, Smith, Fazzari, McNeill, Prestwich and Seymour 2012) raised the possibility that at least in some circumstances such a signal was able to pass from an irradiated fish to a naïve fish in an isolated tank without physical contact between the water in each tank, suggesting the transmission of light or sound. In further experiments reported in the same paper we blocked light signals using aluminium foil and performed attenuation calculations to check if an acoustic signal could be possible. It could not be excluded as the distances and acoustic characteristics of the intervening water and extracellular material are likely to provide efficient sound transmission but was not proven. In a review of the literature in the field we considered the possibility that an acoustic signal might be involved with the initiation or propagation of the bystander signal (Matarèse, Lad, Seymour, Schofield and Mothersill 2020).

It has previously been shown that X-ray photons elicit an acoustic signal from tissues; a property that has been exploited in the development of X-ray-induced acoustic emission tomography (Samant et al. 2020), but these emissions are part of the bulk properties of tissues and the source of emission may be extracellular matrices, water and tissue fluids. The present study is the first to demonstrate an acoustic emission from isolated cells in culture irradiated with X rays at a dose in the range of medical or accidental exposures.

The acoustic signals produced are complex and differ between the two cell types. We are unable to determine the reasons for differences between the emissions from HL-60 and MCF 7 cells but the physical properties of the cells are likely to be the source. The observation that emissions cease almost simultaneously with the cessation of irradiation is consistent with fast physical relaxation processes and not for example the induction of an active mechanism. The hypothesis that RIAE contributes to the bystander effect does not require an energy-dependent process but does require an active sensing mechanism in non-irradiated cells. Such mechanisms have several precedents as discussed in (Matarèse, Lad, Seymour, Schofield and Mothersill 2020), and in some mammalian cell types such as otic hair cells, involve ion (Na^+^, K^+^, Ca^++^) influx and signalling through the action of the MET mechanoreceptor which detects mechanical stretching and deformation of the hair cell membrane (Fettiplace 2017). We consider that it is of significance that calcium influx is the first measurable event in cells receiving a radiation-induced bystander signal (Lyng et al. 2000).

The difference in RIAE pattern from each cell type is likely to be due to the shape, size and elastic properties of the cells. MCF-7 are adherent cells in a monolayer, and HL-60 non-adherent though settled onto the substrate in this experiment. MCF-7 cells have an average interphase size of 15-17μm, while HL-60 cells are smaller at 12.4+-1 μm, and they differ in elastic modulus with HL-60 *E*_a_ = 0.53 kPa. and MCF-7 *E*_a_ = 2.1 kPa. which is a measure of cellular deformability (Rosenbluth et al. 2006). The elasticity and deformability of cells will in principle affect their ability to initiate and respond to an acoustic signal and may affect the nature of the collective pressure wave elicited from each cell type (Nyberg et al. 2017). It is of note that the cell status affects deformability or elasticity, and dead, dying or challenged cells behave differently to normal cells (Otto et al. 2015), suggesting that although the generation of RIAE might be a non-energy dependent process it may still depend on the structure and state of the living cell.

Both HL-60 and MCF 7 cells have previously been shown to generate a bystander effect (Al-Mayah et al. 2012, Pandey et al. 2011a) and we show here that bystander inducing activity is released from these cells in our hands in the same experiment where we measure an acoustic response. HL-60 conditioned irradiated medium elicits a stronger response than MCF-7 in HCT116 recipient (reporter) cells on both cell damage (LDH) and changes in mitochondrial membrane potentials. For these assays the effect of medium from MCF-7 cells show a *p* value of 0.0158, and 0.0357 for LDH and mitochondrial membrane assays respectively and for HL-60 *p*= 0.8, and 0.0357. The very small mitochondrial effect caused in the bystander effect elicited by MCF-7 cells previously reported in microbeam experiments (Shao, Folkard, Held and Prise 2008) may indicate a differential mechanisms in adherent cells in adherent versus non-adherent or cells making intimate cell-cell contacts.

## Conclusion

We demonstrate that under conditions where cultured cells release bystander effect-inducing signals into medium, irradiation of two different cells lines results in the emission of acoustic signals (Radiation Induced Acoustic Emission: RIAE). This phenomenon has not been previously reported for cells in culture, and presents an additional potential mechanism for generation or transmission of the bystander effect that is consistent with unexplained results from other systems. It may also be associated with or mediate the release of other proposed signalling factors such as exosomes. We cannot in these preliminary experiments directly and mechanistically link the bystander effect with acoustic emission, but further work is in progress to establish the relationship between sonic emission and sensing, and the bystander response in vitro. We conclude that irradiation of cultured cells results in a characteristic sonic emission and that this phenomenon needs now to be taken into account when explaining the effects of radiation on biota.

## Acknowledgements and Funding

Work in the laboratory of CM is funded by the Canadian NSERC Discovery Grant 660340. We gratefully acknowledge the excellent technical support of Annette Preston and the generosity of Dr Jane Dobson and the Queens Veterinary School Hospital, Cambridge University Department of Veterinary Medicine for access to the Varian Clinac 2100.

## Notes on contributors

**Dr. Bruno Matarèse** is a physicist in biology and medicine with a particular interest in radiobiology and the use of combined electromagnetic and acoustic technology for early detection of cancer.

**Dr. Hassan Rahmoune** is a cell biologist at the Department of Chemical Engineering & Biotechnology at Cambridge University with particular interest in engineering and biomedical research.

**Dr. Nguyen Vo** was a research associate in the Department of Biology at McMaster University and is currently a research associate in the Department of Biology at the University of Waterloo. He is currently part of a research team mapping the acquired immunity profile for COVID19 in a campus population.

**Prof. Carmel Mothersill** is a radiobiologist with a particular interest in low dose and non-targeted effects of radiation in the environment. She is a professor in the Department of Biology at McMaster University.

**Prof. Paul Schofield** is Professor in Biomedical Informatics at the University of Cambridge and Adjunct Professor at the Jackson Laboratory, Bar Harbor, USA.

**Prof. Colin Seymour** is a radiobiologist with a special interest in low dose effects of radiation. He is a professor in the Department of Biology at McMaster University

## Data availability statement

Raw acoustic data and measurements of extracellular Lactate dehydrogenase and mitochondrial membrane depolarisation are available from the STORE database.

## Data deposition

Data are deposited in STOREDB <u>DOI:10.20348/STOREDB/1169</u>. Individual files can be accessed on:

*STOREDB:DATASET1244* Lactate dehydrogenase assay [<u>DOI:10.20348/STOREDB/1169/1244</u>]

*STOREDB:DATASET1245* Raw Picoscope data [<u>DOI:10.20348/STOREDB/1169/1245</u>]

*STOREDB:DATASET1243* Mitochondrial membrane potential measurements [<u>DOI:10.20348/STOREDB/1169/1243</u>]

